# Climate Change, Fisheries Management, and Increases in Demersal Fish Distribution in a Southern Ocean Biodiversity Hotspot

**DOI:** 10.1101/2025.05.27.656478

**Authors:** Joel Williams, Scott Foster, Skip Woolley, Philippe Ziegler, Cara Masere, Kaitlin Naughton, Otso Ovaskainen, Craig Johnson, Nicole Hill

## Abstract

The world’s oceans and their biodiversity are undergoing going change driven by climate change and anthropogenic impacts such as fishing. The Kerguelen Plateau is a biodiversity hotspot with many endemic fish species, furthermore the region has economic importance in supporting valuable fisheries. This region is also a climate change hotspot with known notable changes in the location of the polar front, ocean currents, and primary productivity. In this study we use data from long-term scientific trawl surveys and contemporary joint species distribution models to understand how the demersal fish assemblage of the Kerguelen Plateau has changed through time and space. The modelling indicates that most demersal species have had notable changes in their occurrence and CPUE from 2003 to 2016. This included a significant increase in species richness throughout the study period. The modelling also provides novel insights into the depth, climatic and environmental preferences for all species, including many species that were previously data limited. It is unclear whether these changes reflect shifts in the fishery, management, or the effects of climate change, but most likely a combination of all. We also found evidence of several species’ distributions responding to temperature variability, with species being exposed to the ongoing impacts of climate change. This new information can be used by managers and policy makers to ensure continuation of sustainable fisheries and the protection of biodiversity into the future.

## 1 | INTRODUCTION

The world’s oceans and their ecosystems are changing due to accumulating anthropogenic pressures such as climate change, extractive activities (fishing, mining etc), and pollution (litter, land runoff, etc.) (Coll et al., 2008; Cooley et al., 2022; Hoegh-Guldberg & Bruno, 2010; Jennings & Kaiser, 1998; Pauly et al., 2005). Over exploitation of a single species or degradation of a habitat can have dire consequence at the ecosystem level (Coll et al., 2008; Jackson et al., 2001; Pauly et al., 1998; Worm et al., 2006). Climate change adds another layer of complexity and uncertainty to the pressures on ecosystems. Global ocean water temperatures are warming or cooling, pH is altering, CO_2_ sequestration is changing, ocean currents are shifting direction and intensity, and the timing and quantity of sea ice melt is changing, all of which is resulting in ecosystem level impacts (Boyd et al., 2016; Constable et al., 2014; Johnson et al., 2011; Wassmann et al., 2011). The changing oceans are resulting in substantial shifts in species’ distributions and relative abundances, often manifesting as range extensions or contractions (Johnson et al., 2011; Schickele et al., 2021). Understanding how anthropogenic pressures and climate effects influence a species or whole communities is important to aid adaptive management, monitoring programs, and identify areas that require protection or greater management to avoid further loss (Arafeh-Dalmau et al., 2021; Emblemsvåg et al., 2022). In our study, we use novel joint species distribution models to examine how the demersal fish assemblage, in a climate change and fisheries hotspot, of the Southern Ocean have changed over recent decades and outline the implications for management in this region.

The Southern Ocean represents approximately 10% of the world’s oceans and it plays a significant role in global primary production, the export of nutrients and oxygen to the world’s oceans, and supports valuable biodiversity (Auger et al., 2021; Constable et al., 2014; Le Quéré et al., 2007; Van de Putte et al., 2021). However, analysis of various properties of the Southern Ocean, such as water temperature, CO_2_ sequestration, and ocean currents reveal that there are large scale changes taking place (Constable et al., 2014; Van de Putte et al., 2021).

The biodiversity of the Southern Ocean is also unique and characterised by a high level of endemism in fish species (Constable et al., 2014), notably the suborder Notothenioids of which 86% of species are endemic to this region (Eastman & McCune, 2000). This group of fishes is also the most abundant on the shelf regions of the Antarctic continent as well as both Antarctic and Subantarctic Islands such as Heard Island and McDonald Islands (HIMI). Temporal changes in Southern Ocean demersal fish composition, distribution, and abundance are mostly unknown (Subramaniam, Melbourne-Thomas, et al., 2020). It is expected that southward shifting ocean frontal systems are likely to have the largest influence on species distributions. However, interpretations of changes in distribution and abundance of fish communities need to be cautious given the confounding effects of fishing and fisheries management as well as other environmental change. Further, the organisation responsible for Commission for the Conservation of Antarctic Marine Living Resources (CCAMLR) have acknowledged the need to better incorporate the effects of climate change into decisions on resource, biodiversity, and ecosystem management in the Southern Ocean (Constable, 2000; Cresswell et al., 2021).

A dominant feature in the Southern Ocean, located halfway between South Africa and Australia, is the Kerguelen Plateau (Duhamel & Welsford, 2011). The Plateau is a productivity hotspot, supporting a diversity of marine life as well as a lucrative and well-managed demersal fishery, primarily for Patagonian toothfish (*Dissostichus eleginoides*) (Duhamel & Welsford, 2011; Hill et al., 2017). Both the location and geography of the Kerguelen Plateau means it is highly exposed to the effects of climate change through warming waters and changing ocean currents and polar fronts (Azarian et al., 2023, 2024). Historically, the region was also exposed to significant illegal, unreported, and unregulated (IUU) fishing. The central part of the plateau surrounding HIMI lies within an Australian EEZ and is managed by Australian Fisheries Management Authority (AFMA) in accordance with CCAMLR ecosystem-based principles. The significance of biodiversity in this region was recognised with the establishment of no-take marine reserves in 2002. Research in this region has focussed on species of commercial and by-catch importance. There are notable gaps in understanding of the composition and structure of fish assemblages along with knowledge about how these communities may have changed through time (Hill et al., 2017). Knowledge and information on how climate change and other factors are influencing the ecosystem is important for the management and conservation of biodiversity and for managers to meet the obligations of the CCAMLR convention. It is also necessary for fisheries’ agencies, such as the AFMA, to ensure adherence to ecosystem-based management practices.

A valuable approach to understanding these environmental processes and ecological change is to explore trends and patterns within robust long-term data sets. Oceanographic data has been collected over large spatial scales for decades (e.g. NOAA). However, long term (>10 years) biological monitoring or ecological datasets (i.e. species’ occurrence and abundance data), while critical for detecting ecological changes, are relatively rare. In the era of big data, increased computing capacity and innovative approaches to ecological modelling, researcher’s abilities to ask more complex questions and to model at larger spatial and temporal scales had become increasingly possible (Franklin et al., 2017; Tikhonov, Duan, et al., 2020). Furthermore, quantitative methods have developed from single species approaches to methodologies which consider whole species assemblage data across large spatial scales using approaches such as joint species distribution models (Ovaskainen et al., 2017; Warton et al., 2015). These modelling approaches allow ecologists to explore correlations across environmental gradients and produce full-coverage ecological maps for each species as well as community attributes such as species richness. These kinds of statistical techniques also enable future predictions under different scenarios (Evans, 2012).

In the study we aimed to assess whether the demersal fish community of the HIMI region of the Kerguelen Plateau has changed through time. We relate the observed changes in presence/absence and abundance of demersal fishes to environmental change, marine reserve zoning, and changes in fishery management practice. To achieve this, we apply joint species distribution modelling (JSDM; (Ovaskainen et al., 2017)). We establish how much of the variation in each species presence/absence and abundance is due to environmental filtering and random process, and how these factors vary across spatial and temporal scales. These models were then used to spatially predict the distribution of each species and species richness for each year of the study. These predictions were compared through time to investigate where and when any change in distribution occurred across the study site. The results from this study provide a useful case study and point of comparison for other regions of the Southern Ocean, which will facilitate better understanding of the likely effects of climate change, fisheries management, and conservation management at a much larger scale.

## 2 | METHODS

### 2.1 | Study site

The management of the Kerguelen Plateau is split between France, who manage the northern half of the Plateau, and Australia, who manage the central portion of the Plateau. This study area was completed within the Australian EEZ on the Kerguelen Plateau in the Subantarctic region of the Indian Ocean (Figure 1). The Plateau itself rises from depths >3000 m, has steep banks, comprises numerous seamounts, and breaks the surface with three large islands and numerous rocky outcrops. The study region is managed and protected by the World Heritage listed Heard Island and McDonald Islands Marine Reserve (Figure 1). During this study this included four areas that are listed as IUCN category 1A marine reserves. The HIMI Marine Reserve was established in 2002, and the reserve boundaries were extended in 2014 to encompass a total of 71,000 km^2^ of ocean. It should be noted that these boundaries were further extend in January 2025 but this is not considered for this study. Sustainable fisheries for Patagonian Toothfish and Mackerel Icefish are also present within the Australian EEZ. The fishery started as a trawl fishery in the 1990s but in 2003 started changing to a predominantly longline fishery to maximise catches of Patagonian Toothfish while minimising bycatch (Welsford et al., 2011). However, there is still some trawling effort that mostly targets Mackerel Icefish. Other species, such as Marbled Rockcod, *Notothenia rossii* and Grey Rockcod *Lepidonotothen squamifrons* were historically targeted by trawling and illegal fishing activities (Duhamel & Williams, 2011), which have been eliminated since the mid-2000s. The region also supports many long-lived, large, and endemic species, such as *Bathyraja irrasa*, that are caught as by-catch.

**Figure 1.**
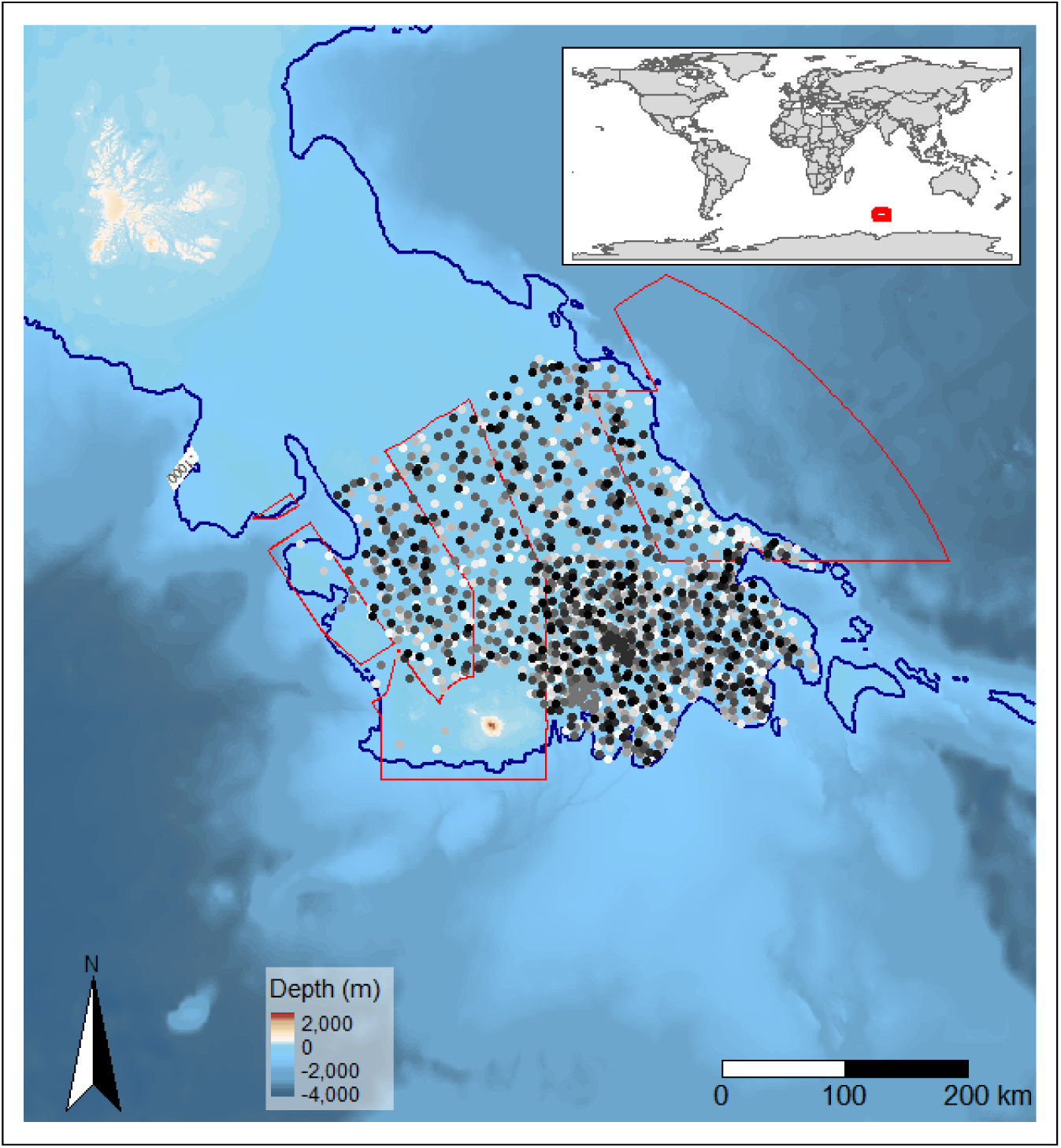
A bathymetric map of the location of each research trawls from the random stratified trawl survey located on the Kerguelen Plateau. The grey-scaled dots represent the year the sample was collected and is scaled 2003 as white through to 2016 as black. The dark blue line represents the 1000m depth contour and the maximum depth of the RSTS samples and limitations spatial predictions in this study. The Heard Island and McDonald Island Marine Reserve boundaries at the time of the study are marked by the red polygons (Zoning changed in 2025). Inset: Location of the study location on the world map.

### 2.2 | Demersal fish data

Demersal fish data used in the study comes from a long-term, annual scientific survey used for monitoring changes in Patagonian Toothfish, mackerel icefish and other bycatch managed species referred to as the random stratified trawl survey (RSTS). The RSTS provides count data for demersal fish in the HIMI region of the Kerguelen Plateau. The RSTS has been conducted each year since 1997, when the fishery began. The survey design divides the HIMI plateau (waters generally less than 1000 m) into 10 strata defined by regions of similar morphology, depth and commercial fishing effort. Sampling sites are randomly located within each of these strata with a consistent number of stations per stratum every year. The survey design and methodology has remained largely unchanged since 1997. This study focused on the period between 2003 and 2016 as this has the best spatial coverage, taxonomic QA/QC, and matching environmental data.

The RSTS uses an otter beam trawl net with a 500 mm mesh cod end liner and is towed at a speed of 3 knots (1.5 m/s) for a duration of 30 minutes. The RSTS occurs from a fishing vessel and trained scientific observers record catches using the same methods as the fishery observer program. Abundance data is available for all species. Tow duration was relatively consistent however variability in environmental conditions provided some differences in trawl distance (min = 1.3 km, max = 6.4 km, mean = 2.9 km, SD = 0.3 km). Given these differences, for the purposes of modelling, the count estimates are divided by the towed distance to provide catch per unit effort (CPUE) (number of fish per swept area).

### 2.3 | Environmental variables

We selected 12 environmental variables that could potentially help explain any change in demersal fish presence/absence, abundance, and community structure (Table 1). This included data derived from the trawls, combined satellite and best available multibeam bathymetry data, satellite sea surface temperature, climate indices, and outputs from oceanographic models. All environmental variables were tested for correlations using Pearsons correlation to ensure pairwise coefficients were <|0.7| to avoid collinearity.

**Table 1.**
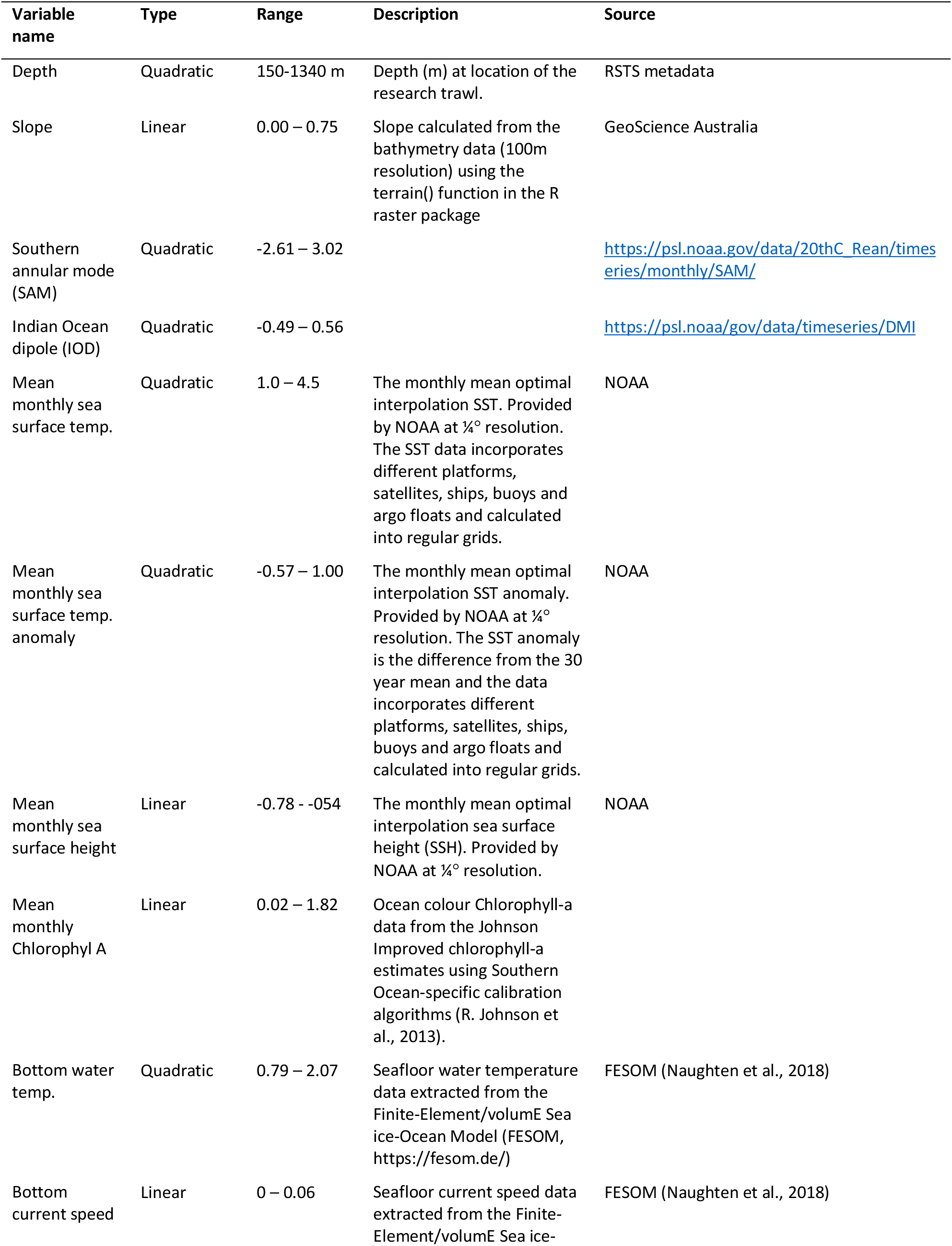
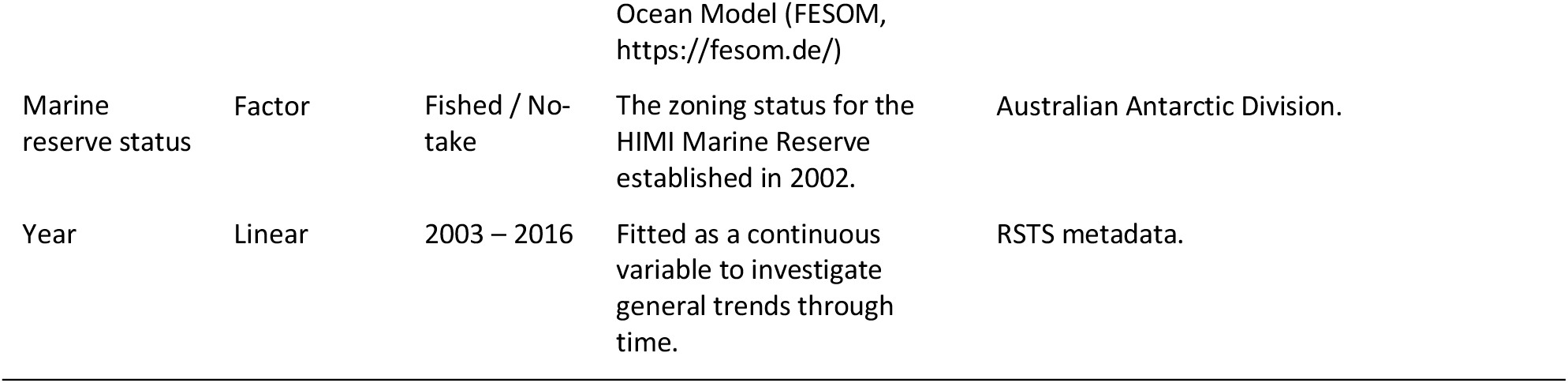
Descriptions of covariates used in the hierarchical modelling of species communities (HMSC).

### 2.4 | Statistical analyses

We analysed the RSTS data using Hierarchical Modelling of Species Communities (HMSC) (Ovaskainen et al., 2017; Ovaskainen & Abrego, 2020). HMSC is a joint species distribution model (Warton et al., 2015) that includes a hierarchical layer to assess the extent to which a species response to environmental covariates depends on species-species and, optionally, species’ traits relationships (Abrego et al., 2017). HMSC also utilizes spatially structured latent variables as proposed by Ovaskainen et al. (2017) and later expanded to big spatial data (i.e. >1000s points or samples) by Tikhonov et al. (2020). The approach enables investigation of how each individual species responds to the environmental covariates, while also considering species co-occurrences through space.

The RSTS data includes records for 38 fish, shark, and skate species sampled from between 111 and 195 trawl sites annually from 2003 to 2016. We excluded those species that had less than 10 occurrences in the data set, resulting in 35 species (*n*_*s*_). The 2,186 individual trawls were treated as sampling units (*n*_*y*_). The response variable was CPUE for each of the 35 species (the matrix *n*_*y*_ × *n*_*s*_ = *Y*). Due to the zero-inflated nature of the data, we applied a hurdle model, i.e. one model for presence/absence and another one for CPUE conditional on presence. We applied probit regression in the presence-absence model, and linear regression for log transformed CPUE data in the CPUE conditional on presence model. The CPUE data were transformed by declaring zeros as missing data, log-transforming the remaining data, and then standardising data within each species (*μ* = 0, *σ* = 1).

We included as fixed effects (the matrix *n*_*y*_ × *n*_*c*_ = **X** of HMSC; see Ovaskainen et al. 2017; where *n*_*c*_ is the number of covariates) 12 variables that could potentially explain the presence and CPUE of the demersal fish assemblage through time (Table 1). We used a sample (trawl) level random factor to account for spatially-structured species’ correlations. Due to the large number of samples (>1,000), and the computational overhead in calculating spatially structured correlation using the default spatial HMSC, we implemented a nearest neighbour Gaussian process using the default 10 neighbours as per Tikhonov et al (2020). Furthermore, due to the study design and the use of sampling strata, we also included stratum as a random factor. To control for potential temporal correlations, we used the numbers of days since the start of the study as a random factor (i.e. 1 January 2003 = 1) for which we assumed exponentially decaying correlation structured with respect to time lag.

We fitted the HMSC model using the R-package Hmsc v3.0-12 (Tikhonov, Opedal, et al., 2020) assuming the default prior distributions (see Chapter 8 of Ovaskainen and Abrego 2020). We sampled the posterior distribution with four Markov chain Monte Carlo (MCMC) chains. The burn-in was set to omit the first 25,000 samples. The four chains were thinned by 100 to yield 500 posterior samples per chain, generating 2000 posterior samples in total. We checked MCMC convergence by ensuring the mean and standard deviation of the potential scale reduction factors were close to unity for the majority of the model parameters (Gelman & Rubin, 1992).

We examined the explanatory and predictive powers of the probit model through species specific area under the curve (AUC) (Pearce & Ferrier, 2000) and Tjur’s R^2^ values (Tjur, 2009). The explanatory and predictive powers of the abundance COP model were measured by R^2^. To compute explanatory power, we used the full dataset to fit models. To compute predictive power, we performed 2-fold cross validation (due to computation time and resources), in which the sampling units were assigned randomly to two folds, and prediction for each fold were based on model fitted to data on the remaining fold.

To quantify the drivers of community structure, we partitioned the explained variation among the fixed and random effects included in the model. To address our main study question, i.e. whether and how species communities have changed over the study period, we examined species responses to the explanatory variables, counting the proportion of species that showed a positive or negative response with at least 95% posterior probability.

Both presence-absence and CPUE conditional of presence models were used to make predictions to produce species distribution maps. A predictive grid was generated at 20×20 km resolution across the study area from 200 m to 1,000 m depth, aligning with the depth limits of the RSTS survey. Probability of occurrence and CPUE conditional on presence was estimated for every grid cell, for every species, for the 1^st^ April every year. Using these predictions, we generated species distribution rasters. To investigate where the changes in distribution were occurring with the study site, we generated maps by taking the mean across the first three years and last three years of the study. This was to account for minor inter-annual variability. We then subtracted the values for the two rasters to establish where change had occurred. Species richness for every year was estimated by summing the predicted occurrence of each species in each cell and changes in species richness was estimated using the same method for individual species.

## 3 | RESULTS

The 35 species included in the modelling comprised of 29 ray-finned fish and six Chondrichthyes. The species assemblage was dominated by three commercially targeted fishery species, *C. gunnari*, *D. eleginoides*, and *Channichthys rhinoceratus*, that represented >70% of the total CPUE across all years. *D. eleginoides* was the most ubiquitous species being present in 91% of trawls, follow by *C. rhinoceratus* that was present in 56% of trawls.

The MCMC convergence of the presence/absence model was satisfactory; the potential scale reduction factors for the β-parameters (that measure the responses of the species to environmental covariates; Ovaskainen et al. 2017) were on average 1.03 (maximum 1.55) for the presence-absence model and for the CPUE conditional on presence model a mean value of 1.00 (maximum 1.01).

### 3.1 | Species responses

The presence/absence model showed a variable fit to the data, the Tjur R^2^ values ranged from 0.06 to 0.78 among species, with the mean being 0.33 (SD 0.22) (Figure 2a), The AUC values ranged from 0.75 to 0.99 among species, with the mean being 0.94 (SD 0.06). The five species or groups with the highest Tjur R^2^, or explanatory power, included in order from highest to lowest, *Channichthys rhinoceratus, Champsocephalus gunnari, Gobionotothen acuta, Muraenolepsis micros,* and *Macrourus* sp. On average, the fixed components of the presence/absence model explained 77% of the explained variance for each species (Figure 2b). Specifically, bathymetry related variables explained on average 33% of the variance across species, satellite derived data 15%, climate related variables 14%, year 9%, seafloor modelled oceanographic variables 4%, and marine reserve zoning 0.5% (Figure 2b).

**Figure 2.**
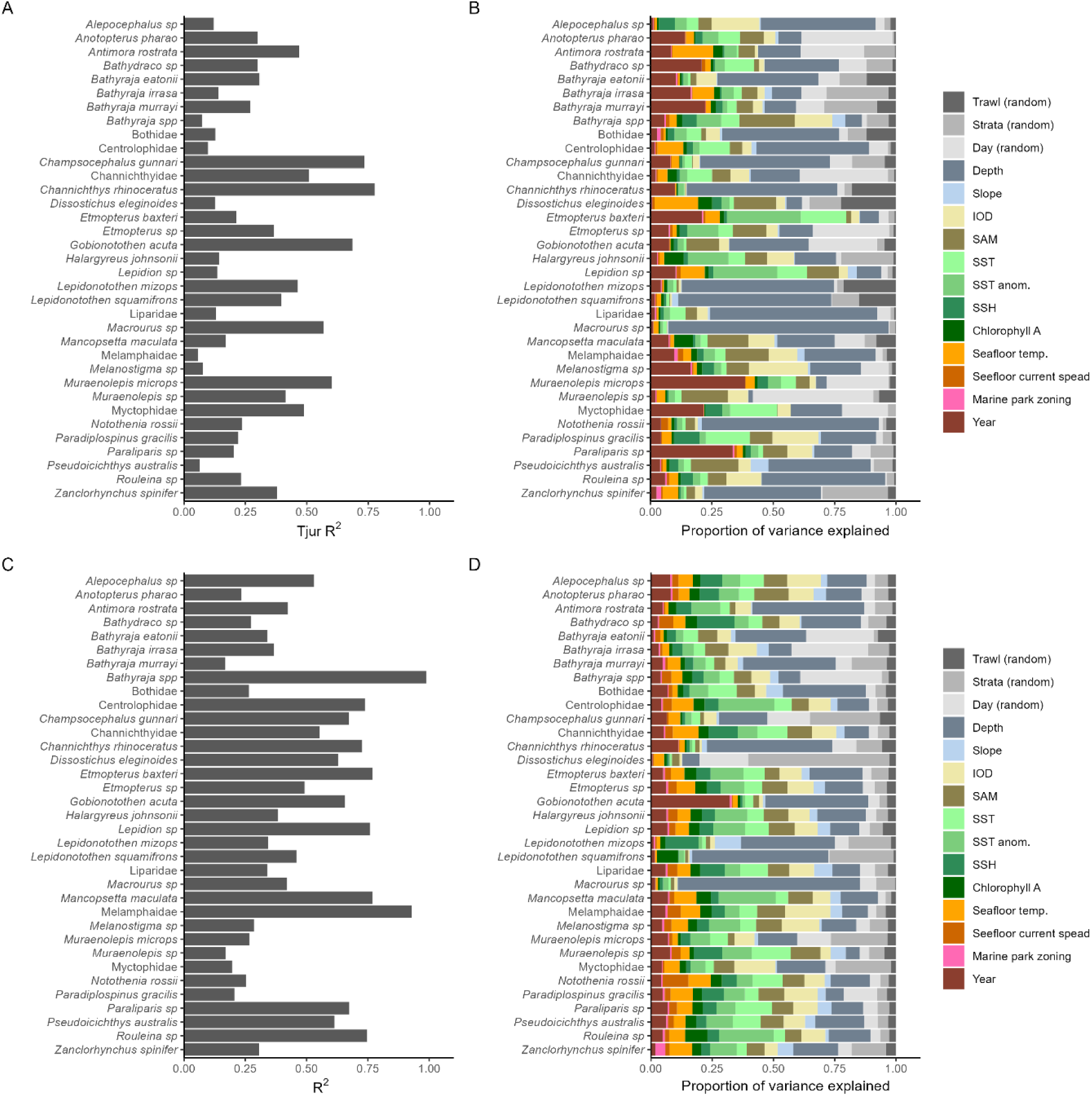
A) The Tjur R^2^ values for each species from the presence/absence model. B) The variation partitioning of the fixed and random effects within the presence/absence model. C) The R^2^ values for each species from the CPUE conditional on presence model. D) The variation partitioning of the fixed and random effects within the CPUE conditional on presence. The species names and species groups are listed in alphabetical order.

The CPUE conditional on presence model showed reasonable fit to the data, the R^2^ values ranges from 0.17 to 0.98 among species, the mean being 0.48 (SD 0.23, Figure 2c). The five species or groups with the highest R^2^, or explanatory power, included in order from highest to lowest are *Bathyraja* sp., Melamphaidae, *Mancopsetta maculate, Etmopterus baxteri,* and *Lepidion* sp. On average the fixed components of the CPUE conditional on presence model explained 79% of the explained variance for each species (Figure 2d). Specifically, bathymetry related variables explained on average 26% of the variance across species, satellite derived data 24%, climate related variables 15%, seafloor modelled oceanographic variables 8%, year 6%, and marine reserve zoning 1% (Figure 2d).

Accounting for only responses that were positive or negative with at least 95% posterior probability, in the presence/absence model 80% of species showed a positive increase in presence across the study period from 2003 to 2016 (Figure 3a). No statistically supported declines in species presence were detected. The CPUE conditional on presence model had 25 % of species showing a statistically supported increase in abundance (Figure 3b). No species were predicted to decline in abundance between 2003 and 2016. Depth and slope were important at explaining the presence/absence of species for 71% and 20% of species respectively showing substantial effect sizes (Figure 3a). Forty-nine percent of species had higher occurrence deeper water, while 23% species in shallow water. Most depth responses also had a significant quadratic second degree term (curved or bell shape gradient) highlighting that many species have well defined depth ranges. For example, *C. gunnari* was only observed in depth <500 m and conversely *Macrourus* sp. was only be seen in depths >500 m (Figure S1). Depth was also most important at describing CPUE conditional on presence with 34% of species having a negative or positive response (Figure 3b). Seafloor slope was significant for 11% of species (Figure 3b).

**Figure 3.**
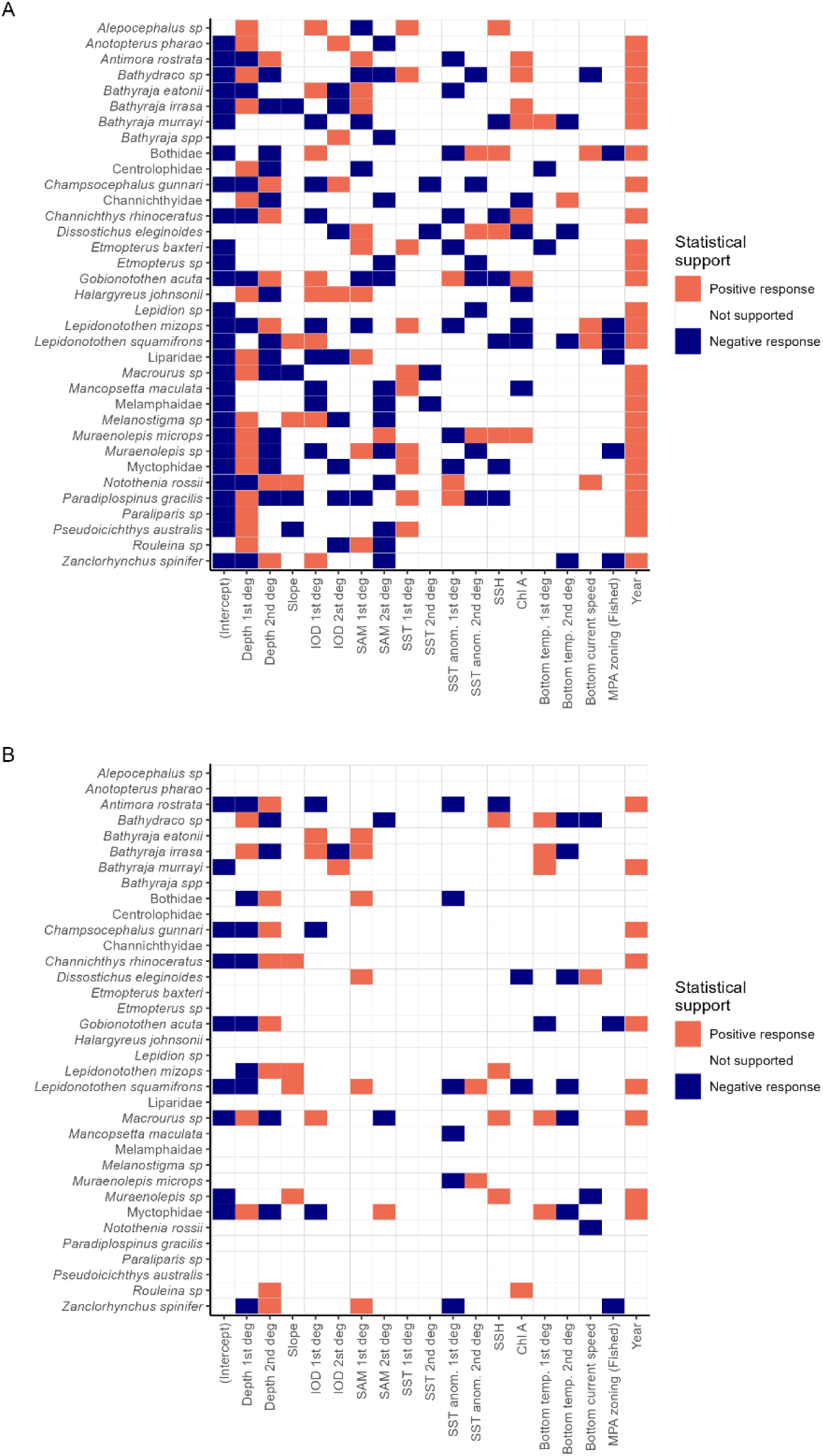
The responses of the species to environmental covariates. Panel A showed the results for the presence-absence model and panel B for the CPUE conditional on presence model. In both panels, responses that are positive with at least 95 % posterior probability are shown in red, response that are negative with at least 95 % posterior probability are shown in blue, and responses that were not statistically supported are shown in white. Note: All variables are continuous except MPA Zoning, which is a 2-level factor with this figure showing the contrast of “Fished” sites when compared to sites in the no-take Marine Reserve. The species and species groups are ordered alphabetically.

The climate variables, IOD and SAM had variable response on the presence of demersal fish species (Figure 3). Almost half of the species included in the presence/absence model had a statistically supported positive or negative response to IOD or SAM (Figure 3a). For CPUE conditional on presence the effect was not as strong with only six species (three positive, three negative) responding to IOD and six species positively responding to SAM (Figure 3b). One third of the species had a positive relationship with SST for presence/absence (Figure 3a). However, there were no statistically supported responses in the CPUE conditional on presence model (Figure 3b). SST anomaly had a mixed response for the presence absence model with three species having a significant positive relationship and eight species having a negative relationship (Figure 3a). SSH, Chlorophyl A, and the seafloor modelled variables had a notably smaller number of species with a statistically supported responses for the presence/absence model (Figure 3b).

The presence of six demersal fish species was higher inside the no-take marine reserve when compared to fished areas, including two fishery managed by-species *Lepidonotothen mizops* and *L. squamifrons* (Figure 3a). For the abundance COP model the only species to show a statistically supported response were *Gobionotohen acuta* and *Zanchlorhynchus spinifer* that were more abundant inside the no-take marine reserve (Figure 3b).

### 3.2 | Changes in species distribution

There was statistical support for an increase in the presence of 80% of the species modelled across the duration of this study (Figure 3). Spatial predictions converted to rasters demonstrate the spatial extent of these changes (Figure 4). Changes in occurrences were highly species specific (Figure 4). Maps for all three skate species showed similar changes in occurrence with 20-30% increases in presence across the top of the plateau in low relief areas. The change in occurrence for *B. eatonii* (Figure 4) were more widespread than for *B. murrayi* and *B. irrasa*. The distribution of the fisheries targeted *C. gunnari* was more concentrated to the shallower areas of the Plateau including to the north of Heard Island, and on top of ridges and rises (Figure 4). The greatest increases in occurrence were to the north of Heard Island. *L. squamifrons* was distributed across the Plateau, with the greatest change occurring to the northern extent of the Australian EEZ (Figure 4). In contrast, *Macrourus* sp. tended to be more associated with deeper and high relief section of the Plateau with the greatest increases in occurrence on slope to the south and north (Figure 4).

**Figure 4.**
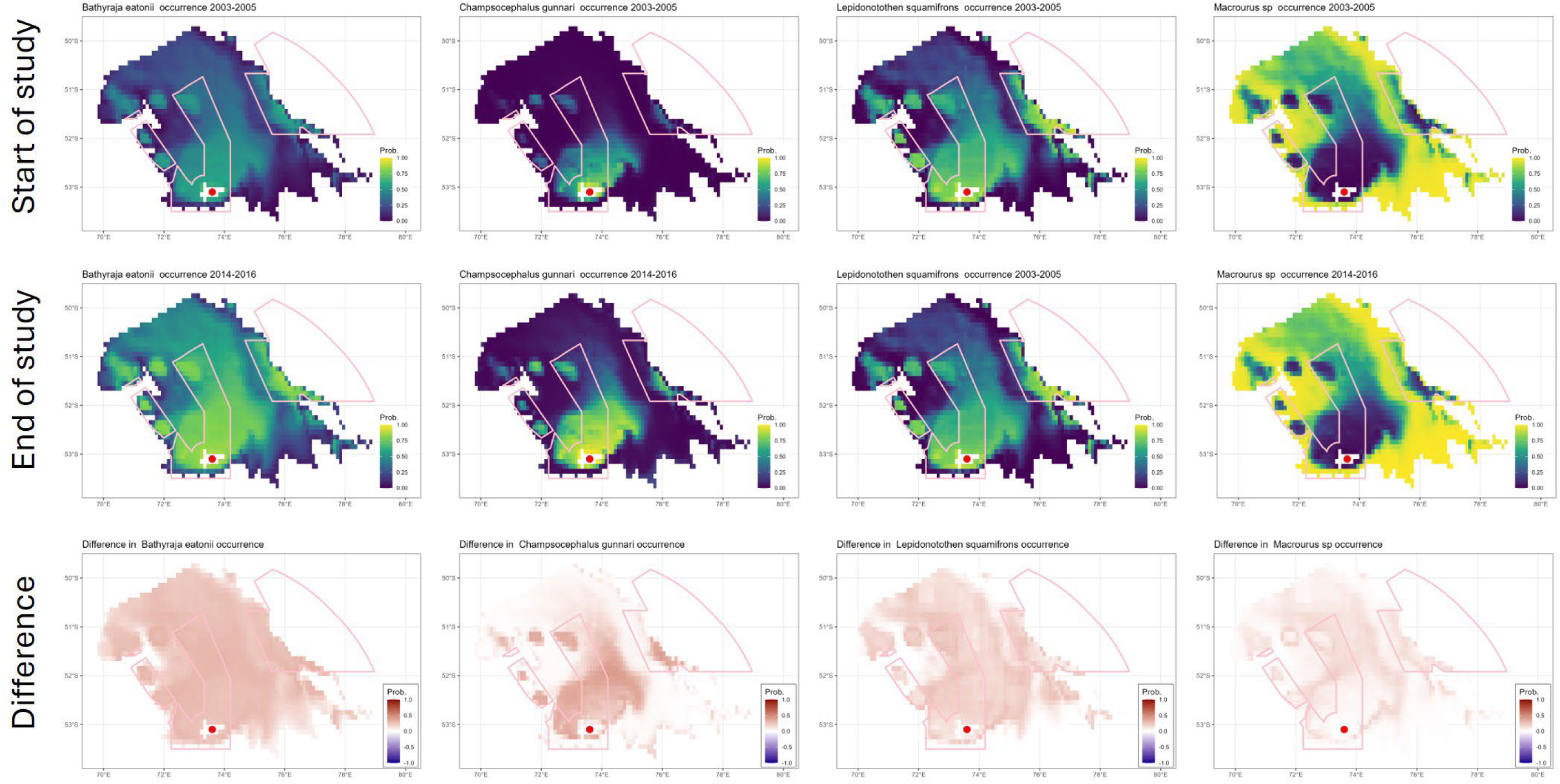
Change in distribution of the four species of interest, taking the mean probability of occurrence from the first three years (2003-2005) and the last three years (2014-2016) and calculating the difference in probability of occurrence between the two period. For reference, the red dot represents the location of Heard Island, and the pink polygon represent the HIMI Marine Reserve boundaries at the time of the study.

There was statistical support for an increase in CPUE conditional of presence for 30% of species modelled. The maps demonstrate that if the species is present then this is how abundant it is at the location and time. Despite a significant increase in the occurrence of *B. eatoniii*, there was only a very small increase in CPUE with a maximum increase of 0.3 individuals CPUE (Figure 5). This was on the shallower and low relief sections of the Plateau (Figure 5). In contrast, *C. gunneri* show increases in CPUE around Heard Island and along a ridge to the east (Figure 5). CPUE of *L. squamifrons* showed similar patterns with a notable increase in CPUE along a ridge to the east (Figure 5). Macrourus sp. had notable increase in CPUE across the deeper slope regions of the Plateau (Figure 5).

**Figure 5.**
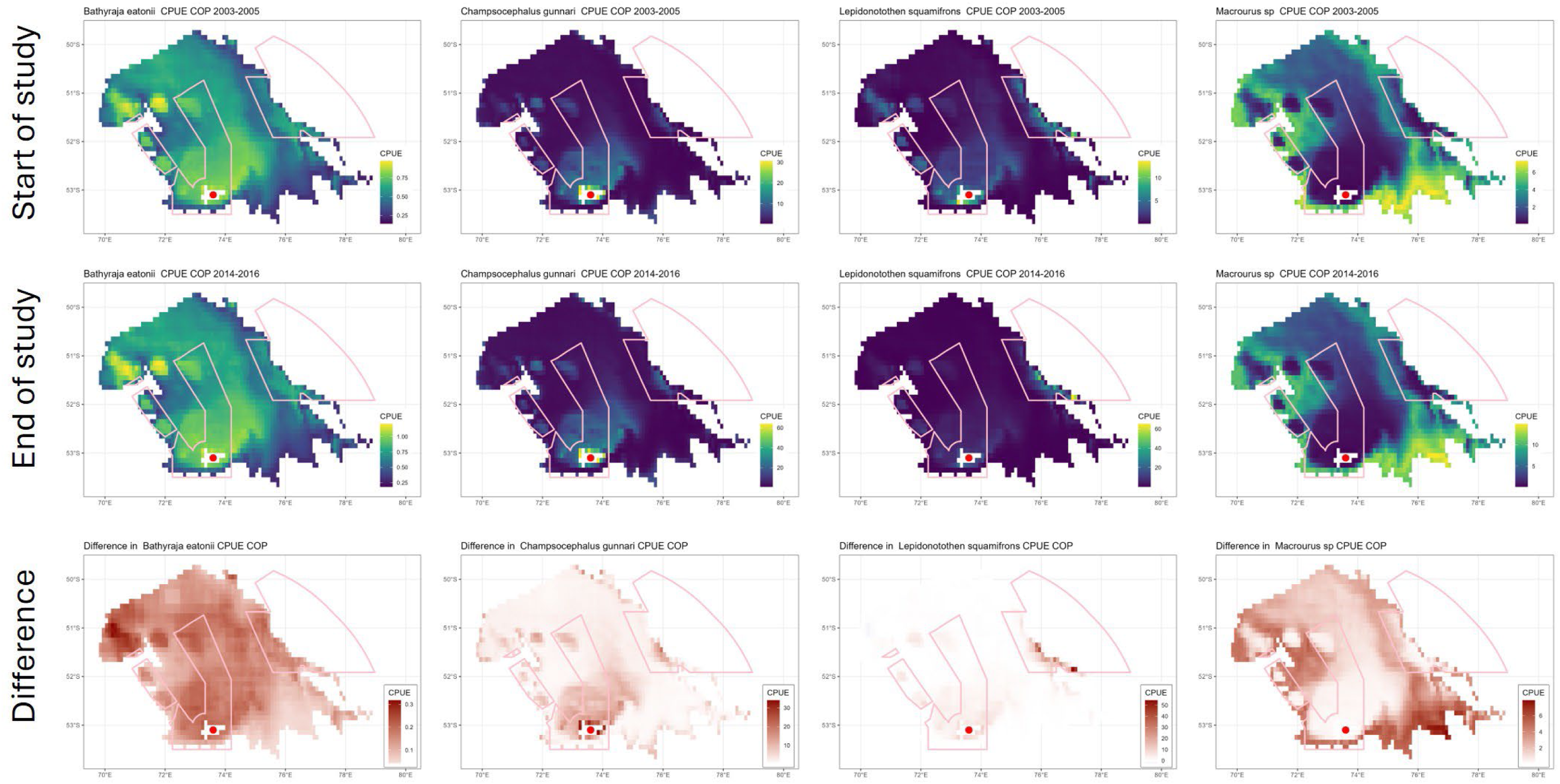
Change in distribution of CPUE conditional of presence from the first three years (2003-2005) and the last three years (2014-2016) and calculating the difference in probability of CPUE conditional of presence between the two period. For reference, the red dot represents the location of Heard Island and the pink polygon represent the HIMI Marine Reserve boundaries at the time of this study.

### 3.2 | Species richness

At the assemblage level, the HMSC model predicted that species richness increased through time, with a mean species richness of three species per trawl in 2003 increasing to seven species per trawl in 2016 (Figure 5). Species richness also decreased with depth from a mean of eight species per trawl in 200 m depth to five species per trawl in 1,000 m depth (Figure 6). There was a positive correlation with species richness and SST, with species richness more than doubling within an increase of 3°C of sea surface temperature (Figure 5).

**Figure 5.**
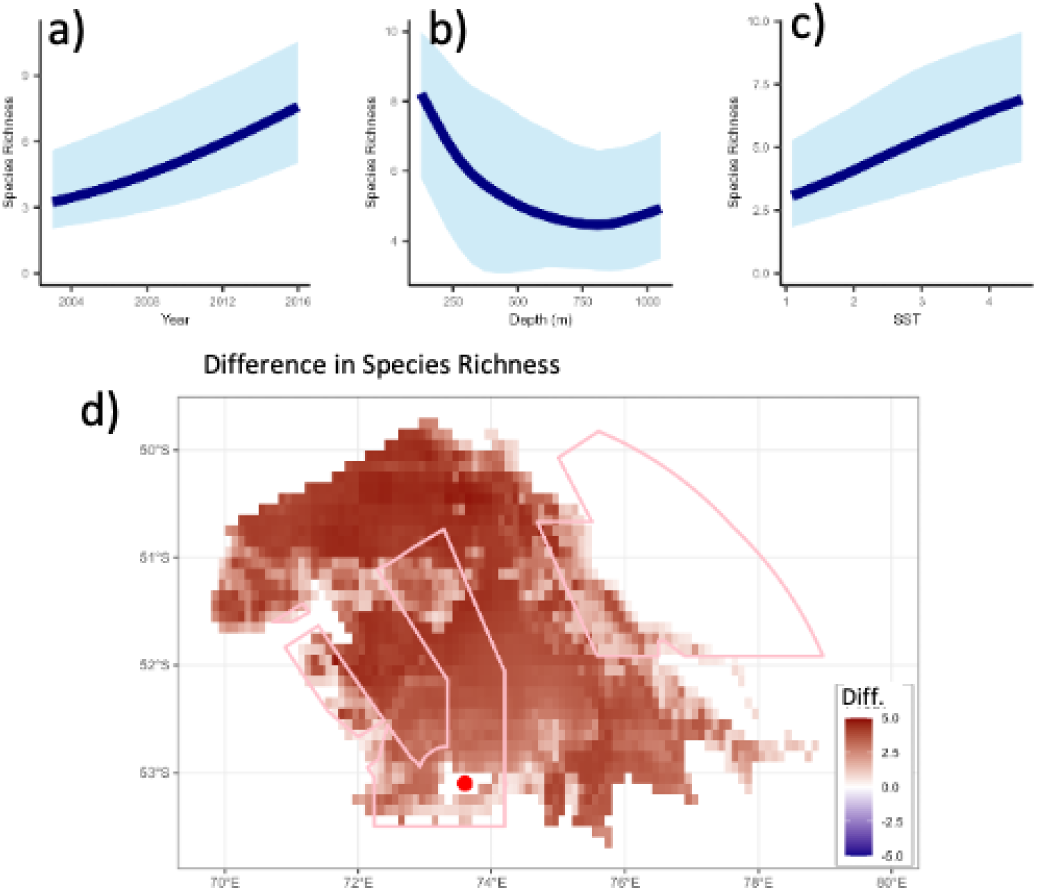
Model predictions of species richness through (a) time (Year), (b) across depth, and across(c) sea surface temperature (SST). (d)The difference in species richness between the first three year (2003-2005) and the last three year (2014-2016). The pink polygons represent the HIMI Marine Reserve boundaries at the time of the study, and the red dot is the location of Heard Island.

## 4 | Discussion

In this study we have demonstrated how the demersal fish community in the Heard Island and McDonald Island (HIMI) region of the Kerguelen Plateau has changed from 2003 to 2016. We detected increases in species occurrence, CPUE, and species richness. This study has similar findings and draws the same conclusions as a study from the French EEZ on the Kerguelen Plateau, noting with much fewer samples (Duhamel et al., 2019). It is well documented that the Kerguelen Plateau ocean climate has also changed and will continue to change into the future (Cavanagh et al., 2021; Pinkerton et al., 2021; Su et al., 2021). Furthermore, over the last few decades, management practices have improved, there has been an elimination of illegal, unreported, unregulated (IUU) fishing, combined with fishery-led changes to minimise by-catch. This multitude of factors has likely contributed to a cumulative influence on the demersal fish community. While, the structure of the species assemblage can be defined largely by physical geography, i.e. depth and slope, there is evidence that some change in the fish community can be attributed to shifts in climate and oceanography. At the species level, the targeted fishery species, Patagonian Toothfish were ubiquitous in distribution across the study area and throughout the study period. In comparison Mackerel Icefish were concentrated around particular areas to the north and east of Heard Island and across the shelf edge on the east of the Plateau. It is the fishery by-catch species, i.e. non-targeted species, that are not studied and yet potentially sensitive to fishery pressures, and it is these species that have shown increases in both prevalence and abundance in this study. This suggests that any or all the elimination of IUU fishing, improved fisheries management, and shifts in fishing methods have all had a positive influence on these species.

The advantage of having long-term data used with joint species distribution modelling has proven vital in providing novel insights into individual species as well as the fish assemblage and diversity metrics like species richness. These findings are important to understanding the ecology of the Southern Oceana and Antarctic ecosystems.

### 4.1 | Changes in distribution of demersal fish and ecosystem effects

We detected notable changes in the distribution and CPUE of many species in the north of the study site and particularly on banks and ridges. It is well document that the distribution of demersal fish is closely linked to depth, habitat type and structure (Bax et al., 1999; Brown et al., 2022; Paxton et al., 2017). Highly complex habitats support greater biodiversity and increased biomass of demersal fishes (A. F. Johnson et al., 2013; Trebilco et al., 2015). Habitat surveys and modelling of habitats in the HIMI region have shown the highest biodiversity and most complex habitats occur on top of banks, rises, and ridges (Welsford et al., 2013). It is plausible that with the reduction in trawl effect, that there has been recovery of habitat in these regions and that could explain some these changes in the demersal fish assemblage.

Demersal fish are prey for mammals and birds highlighting their importance in food-web and ecosystem dynamics (McCormack et al., 2021). The are several ecosystem models for the region but they tend to group fish into broad pelagic or demersal groups (Subramaniam, Corney, et al., 2020). As demonstrated in this study demersal fishes often have geographic and environmental niches that aren’t captured at the coarse scale of these ecosystem models. Most demersal species recorded during this study feed on mesopelagic fish species. Mesopelagic species are known to aggregate around areas of primary and secondary productivity (Woods et al., 2023). Bioregionalisation research in this region has found the Plateau to be a region of high productivity supporting a distinct group of zooplankton (Chapman et al., 2020; Godet et al., 2020). However, with zooplankton being closely linked to ocean conditions, climate change is likely to be influencing primary productivity that is likely to flow through the food chain and influence demersal fishes (A. J. Constable et al., 2014; Godet et al., 2020; Richardson, 2008; Su et al., 2021).

### 4.2 | Changes related to climate and oceanography

It is also feasible that changes in environmental conditions, for example higher SST, have increased productivity to the benefit of demersal fishes. Warming waters in high latitude areas has been linked to an increase in primary productivity due to reduced water column mixing (Doney, 2006).

Chlorophyll-a concentration (from satellite data) showed mixed results for prevalence and little explanation for abundance (Doney, 2006). One of our hypotheses, that need further exploring, is that changes to ocean currents and climate have led to increase in overall productivity and this is flowing through to an increase in the demersal fish community. The Kerguelen Plateau is the largest topographic feature and barrier to the eastward moving Antarctic Circumpolar Current (ACC; (van Wijk et al., 2010). Therefore, the region has complex oceanography with the polar and sub-polar fronts converging on the northern slopes of the Plateau and significant currents passing through the troughs between Kerguelen island and Heard Island, and between Heard Island and Banzare Bank (van Wijk et al., 2010; Welsford et al., 2013). This is one of the reasons why this region has high productivity and unique and important biodiversity values. It also means the region is particularly susceptible to climate change driven changes to ocean currents and climate. Climate change is already influencing Southern Ocean and there is evidence of the polar fronts moving southward and that long-term gradual warming is already occurring at depth (Gille, 2002; Haumann et al., 2020). Extreme pulse events such as marine heatwaves are impacting this region (Azarian et al., 2023; Rogers et al., 2015; Su et al., 2021). There is also growing evidence of cold water events. It is unclear how these extreme events such are impacting demersal communities in remote areas of the Southern Ocean. There is some evidence that suggests Macrourids and *L. mizops* have defined thermal niches. Thus, extreme events such as marine heatwaves are likely to thermally stress these species resulting in physiological impacts or even range or depth shifts or contractions.

This study also found the Southern Annular Mode (SAM) to be a significant climate variable that explains the prevalence and abundance of demersal fish. The response to SAM was mixed and species dependent. The prevalence of some species was highest when SAM was neutral, while other species showed greatest prevalence during positive SAM periods. Positive SAM events have been attributed to high climate variability, poleward shift in front, strengthening westerly winds and marine heatwaves (Fogt & Marshall, 2020; Le Quéré et al., 2007; Su et al., 2021). SAM is believed to be the leading mode of climate variability in this region yet the direct links between SAM and the ecosystem function and dynamics are still relatively unknown (Fogt & Marshall, 2020).

### 4.3 | Changes related to fisheries and spatial management

Two of the most ubiquitous species were the Patagonian toothfish and mackerel icefish fisheries, which are highly managed fisheries. Quotas for fishing within the Australian EEZ and HIMI region are set and reviewed annually by Australian Fisheries Management Authority (AFMA) and the Convention for the Conservation of Antarctic Marine Living Resources (CCAMLR). This includes bycatch limits and other management rules such as ‘move on rules’. The adaptive ecosystem-based management approach applied by AFMA and CCAMLR can, but is yet to, incorporate natural and climate-driven fluctuations in the distribution and stock structure of fishery managed species.

In 2003, the HIMI commercial Patagonian Toothfish fishery started moving away from demersal trawling to long-line fishing (Ziegler & Welsford, 2019). This also included a move to targeting fish in deeper water (>1,000 m). However, it should be noted that some trawling targeting the mackerel icefish still occurs in the shallower region. This change in gear type and locations of fishing was driven by the fishery itself to maximise efficiency and returns and minimise bycatch and benthic impacts. It is undoubtable that seabed trawling has an impact on demersal habitats and recovery times from trawl impacts can be varied from years to decades (Pitcher et al., 2015, 2022). It is plausible that the reduction of trawl fishing effort has allowed for the recovery of habitats to the benefit of demersal fishes. This is coupled with that fact that one of the benefits of long line fishing is that it greatly reduces by-catch and fewer non-targeted species are caught. Similar changes in fisheries management and distribution of fishes has been detected to the north in the French EEZ (Duhamel et al., 2019).

The HIMI region is also spatially managed through no-take marine reserves. The Heard Islan⍰and McDonald Island Marine Reserve was established in 2002 to protect the conservation value and the vulnerable marine ecosystem. In January 2025, the marine reserve was extended to cover nearly 90% of Australia’s EEZ with different zoning types, including extensive areas of no-take zoning (IUCN II). The RSTS, and thus this study, does have samples from within the no-take marine reserve prior to the 2025 extension. However, there is no ‘before’ data and there are few samples per year collected within the original no-take zones. Marine reserve status was included in the modelling, and we did detect no-take zone effects for six species. This included the two Lepidonotothen species, with *L. squamifrons* being a by-catch managed species. The other four species or Family groups included, Bothidae, Liparidae, *Muraenolepis* sp., and *Zanclorhynchus spinifer*. However, it is difficult to confirm if these increases in prevalence and abundance are due to complete protection or if these protected areas already consisted of preferable habitat for this species. More targeted monitoring of no-take marine reserves would be required to establish no-take effects. This RSTS program and data has become more valuable with the recent significant expansion of the marine reserve in January 2025. This means there is now over two decades of before data from regions on the marine reserve in 200-1000m of water. While the re-zoning still allows fishing activities there are new no-take zones that overlap with the spatial coverage of the RSTS program.

### 4.4 | Long-term monitoring to detect change

Overall, this study demonstrates the immense benefit and value of the annual RSTS as a fisheries independent survey. Without long-term data collected using the same comparable methodology it would not be possible to detect these changes accurately and confidently. The random stratified component ensures adequate special coverage and avoids sampling biases that increase or decrease effect sizes. It should be noted that trawl surveys do target lower complexity habitat to minimise risk of entanglement of nets. Therefore, the RSTS is likely to have a bias towards species that inhabit soft sediments and low relief reefs and provide an overall underestimate of abundance and species richness. Other methods, such as long-line surveys, remote underwater video, eDNA are needed to capture baseline data for demersal fish that inhabit the more complex reefs systems that are known to occur in this region. Additional research such as taxonomic and biological (age at length, diet etc) studies should be linked with these surveys. There is also a benefit to expand surveys to deeper waters to capture all life stages and allow for greater spatial coverage.

### 4.5 | Conclusion

This study has demonstrated that demersal fish community has changed over a 13-year period. The prevalence, abundance, and species richness have all increased across the HIMI region of the Kerguelen Plateau. This includes for many of the fisheries bycatch managed species such as the skates and macrourids. While it is difficult to disentangle what has driven this change in distribution and abundance, we conclude that it is likely a combination of factors including improved fisheries management and fishing practices, elimination of illegal fishing, and environmental change because of climate change. For most species this is the first investigation into their distribution through space and time in the HIMI region, providing novel insights into these demersal fish species. Overall, this is good news for the Kerguelen Plateau indicating that currently ecosystem-based management is working. The HIMI fisheries management is built on the ecosystem-based management principles of CCAMLR. To date most of the demersal fish knowledge in the HIMI region has been based on the two fishery species. This study provides valuable data and knowledge the on the demersal fish community that can help with the assessment of fisheries management practices such as bycatch limits and ‘move-on’ rules. Furthermore, the availability of long-term data and the approach using joint species distribution modelling has proved to be highly beneficial for understanding the distribution of demersal fish across the HIMI region.

## Acknowledgements

We are grateful to the Australian fishing industry for providing survey support for the collection of RSTS data. We acknowledge the fisheries observers, crew and skippers for undertaking data collection and reporting, and to T. Lamb for data management and preparation. This work was made possible through an Australian Antarctic Program grant (Project 4501).

## Data Availability Statement

Due to the confidentiality of the benthic fish data, the raw abundance data needs to be requested from the Australian Antarctic Data Centre (https://data.aad.gov.au/metadata/records/HIMI_RSTS_Strata). The environmental data and R code used to wrangle, model the data, and produce predictive maps are open access and available from the Institute for Marine Antarctic Studies Metadata Catalogue (https://doi.org/10.25959/4GVK-RM21).

